# Intraocular pressure elevation precedes a phagocytosis decline in a model of pigmentary glaucoma

**DOI:** 10.1101/175695

**Authors:** Yalong Dang, Susannah Waxman, Chao Wang, Ralitsa T. Loewen, Nils A. Loewen

## Abstract

**Purpose:** Outflow regulation and phagocytosis are key functions of the trabecular meshwork (TM), but it is not clear how the two are related in secondary open angle glaucomas characterized by an increased particle load. We hypothesized that diminished TM phagocytosis is not the primary cause of early ocular hypertension and recreated pigment dispersion in a porcine ex vivo model.

**Materials and Methods:** Sixteen porcine anterior chamber cultures received a continuous infusion of pigment granules (P), while 16 additional anterior chambers served as controls (C). Pressure transducers recorded the intraocular pressure (IOP). The phagocytic capacity of the trabecular meshwork was determined by fluorescent microspheres.

**Results:** The baseline IOPs in P and C were similar (*P*=0.82). A significant IOP elevation occurred in P at 48, 120, and 180 hours (all *P*<0.01, compared to baseline). The pigment did not cause a reduction in TM phagocytosis at 48 hours, when the earliest IOP elevation occurred, but at 120 hours onward (*P*=0.001 compared to C). This reduction did not result in an additional IOP increase at 120 or 180 hours compared to the first IOP elevation at 48 hours (*P*>0.05).

**Conclusion:** In this porcine model of pigmentary glaucoma, an IOP elevation occurs much earlier than when phagocytosis fails, suggesting that two separate mechanisms might be at work.

## Introduction

Conventional outflow can account for up to 85% of total aqueous drainage^1^ and is guarded by the trabecular meshwork (TM), a strainer - like tissue with cells surrounded by variable amounts of extracellular matrix (ECM).^2^ These cells allow the aqueous to pass into Schlemm's canal by paracytosis and directly through the cells by giant vacuoles as the primary mechanism.^3^ A failure to maintain a normal cytoskeleton and homeostasis of aqueous outflow can cause ocular hypertension.^2^ For instance, pigment dispersion^4^ and corticosteroids can alter the actin cytoskeleton and cause TM cell contraction resulting in an elevation of intraocular pressure (IOP).^4,5^ Conversely, relaxing the cytoskeleton, for instance by using a Rho kinase inhibitor, can reverse these effects.^6,7^

Phagocytosis of debris is another key function of TM cells.^3^ However, its direct and short - term effects on intraocular pressure (IOP) regulation remain poorly understood^2^ although chronic exposure to pigment,^8^ erythrocyte - derived ghost cells,^9^ inflammatory cells,^10^ photoreceptor outer segments,^11^ lens and pseudoexfoliation material^12,13^, can result in challenging secondary glaucomas.

We recently developed an ex vivo pigmentary glaucoma (PG) model that recreates the IOP elevation, stress fiber formation, and overwhelmed phagocytosis that is characteristic for human PG.^4^ We observed a 35.3% reduction in TM phagocytosis and creation of long, thick actin stress fibers. A gene expression analysis indicated an activation of the RhoA signaling pathway and a downstream effect of tight junction formation that was negatively regulated by RhoA - mediated actin cytoskeletal reorganization.^4^ In the current study, we hypothesized that ocular hypertension is the result of a reorganization of the actin cytoskeleton and occurs upstream and earlier than the reduced phagocytosis we saw.

## Materials and methods

### Pig eye perfusion culture and pigmentary glaucoma model

This study was conducted in accordance with the Association for Research in Vision and Ophthalmology Statement for the Use of Animals in Ophthalmic and Vision Research. Because no live vertebrate animals were used and pig eyes were acquired from a local abattoir (Thoma Meat Market, Saxonburg, PA), no Institutional Animal Care and Use approval was required. Thirty - two porcine eyes were cultured within 2 hours of enucleation. Extraocular tissues were removed, and the eyes were decontaminated with 5% povidone - iodine solution (CAT# 3955 - 16, United States Pharmacopeia, Rockville, MD) for two minutes and washed three times in phosphate buffered saline (PBS). Posterior segments, lenses, and irises were removed and the anterior segments with intact TM mounted in the perfusion system as previously described.^4,14,15^ We used the same method to generate pigment granules as recently described in a model of pigmentary glaucoma (PG).^4^ Briefly, pigment granules were produced by subjecting the iris to freeze - thaw and resuspension washing before dilution of the stock to a final concentration of 1.67 × 10^7^ particles / ml. Eyes in the pigment dispersion group were continuously perfused with pigment added to the culture medium for up to 180 hours (P) and compared to controls (C). The perfusate consisted of Dulbecco's modified Eagle media (DMEM, SH30284, HyClone, GE Healthcare, UK) that was supplemented with 1% FBS and 1% antibiotics (15240062, Thermo Fisher Scientific, Waltham, MA) at a constant rate of 3 μl/min using a microinfusion pump (PHD 22 / 2000; Harvard Apparatus, Holliston, MA). IOP was measured intracamerally by a pressure transducer (SP844; MEMSCAP, Skoppum, Norway) and recorded at two - minute intervals (LabChart, ADInstruments, Colorado Springs, CO). Baseline IOPs were obtained after IOP stabilization for 48 hours.

### In situ measurement of TM phagocytosis

A suspension of 0.5 μm carboxylate - modified yellow - green fluorescent microspheres^16^ (cat# F8813, Thermo Fisher, Waltham, MA) at 5 × 10^8^ particles / ml was added to the perfusate at 48, 120, and 180 hours and perfused for 24 hours. The eyes were removed from their perfusion dishes, washed three times with pre - warmed PBS, secured again in the perfusion dishes, and placed upside down for imaging. The TM, visualized from the underside of the transparent perfusion dish, was photographed and measured by acquiring the images with a camera and epifluorescence equipped dissecting fluorescence microscope (SZ X 16, Olympus, Tokyo, Japan) at a 680 × 510 pixel resolution and a 200 ms exposure. The mean fluorescence intensity was quantified by ImageJ (Version 1.50i, NIH) as previously described^17^ at 48, 120, and 180 hours by measuring the fluorescence intensity in the TM.

### Histology

After the TM phagocytosis assay, anterior segments were fixed with 4% PFA for 24 hours, washed three times with PBS, dehydrated in 70% ethanol, and embedded in paraffin. Sections were cut to a thickness of 5 μm and stained with hematoxylin and eosin (H&E).

### Statistics

Data were presented as the mean ± standard error and analyzed by PASW Statistics 18 (SPSS Inc., Chicago, IL). The baseline IOP was compared to the other time points of the same eye using a paired *t*-test. Other quantitative data were analyzed by one-way ANOVA. A *p* value ≤ 0.05 was considered statistically significant.

## Results

In H&E stained tissue sections, normal TM (**Fig. 1 A**) presented as a sparsely pigmented (**red arrowheads**), multilayered, porous tissue with Schlemm's canal - like segments within the aqueous plexus at the outer layer (**black arrows**). Pigment perfusion resulted in abundant pigmented granules phagocytosed by meshwork cells, particularly in the uveal TM, at 48, 120, and 180 hours (**Fig. 1 B, C and D**). Pigment granules were not physically obstructing any part of the conventional outflow system. Baseline IOP in P was comparable to that of C (12.2 ± 0.9 mmHg vs. 11.9 ± 0.9 mmHg, *P*=0.82). Pigment dispersion caused a significant IOP elevation at 48, 120, and 180 hours (19.5 ± 1.4 mmHg, 20.2 ± 1.4 mmHg and 22.8 ± 0.8 mmHg, *P*=0.001, P < 0.001 and *P*=0.002, compared to baseline) while IOPs in C remained steady (13.1 ± 1.1 mmHg, 12.0 ± 0.9 mmHg and 14.0 ± 1.5 mmHg, all p values >0.05, compared to baseline) (**Fig. 2 A**). By inverting the perfusion dishes and washing away the microspheres in the intertrabecular spaces, the TM phagocytosis was visualized and quantified under an upright dissecting fluorescence microscope. Pigment did not cause any change of phagocytosis during early ocular hypertension at 48 hours (**Fig 2 B i-ii**, 96.3 ± 5.0% compared to the control, *P*=0.723), but did cause a reduction at the later phases of 120 hours (**Fig 2 B iii-iv**, 58.3 ± 2.3%, *P*=0.001) and 180 hours (**Fig 2 B v-vi**, 62.5 ± 5.1%, *P*=0.026). However, the declining phagocytosis did not result in further elevation of IOP at 120 and 180 hours compared to the initial IOP elevation at 48 hours (20.2 ± 1.4 mmHg and 22.8 ± 0.8 mmHg versus 19.5 ± 1.4 mmHg, both *P*>0.05).

**Figure 1.**
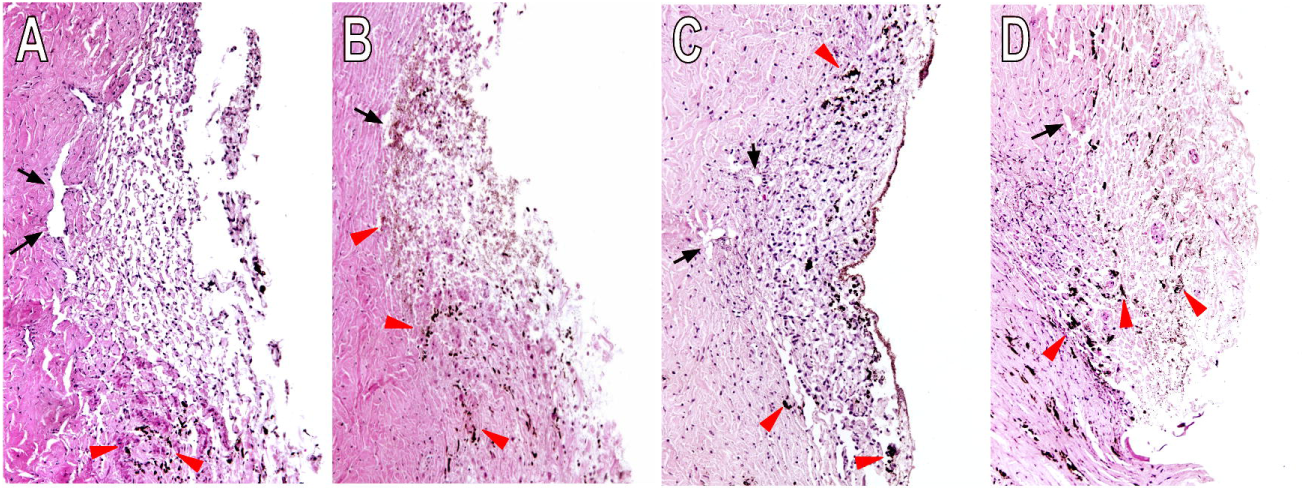
Histology. Normal trabecular meshwork (TM) (**Fig. 1 A**) was a multilayer, strainer - like structure with few pigment deposits (red arrowheads). Ex vivo perfusion with pigment granules at 1.67 × 10^7^/ml caused significant TM pigmentation at 48 hours (**Fig. 1 B**), 120 hours (**Fig. 1 C**) and 180 hours (**Fig. 1 D**). No apparent occlusion to the outflow tract was found.

**Figure 2.**
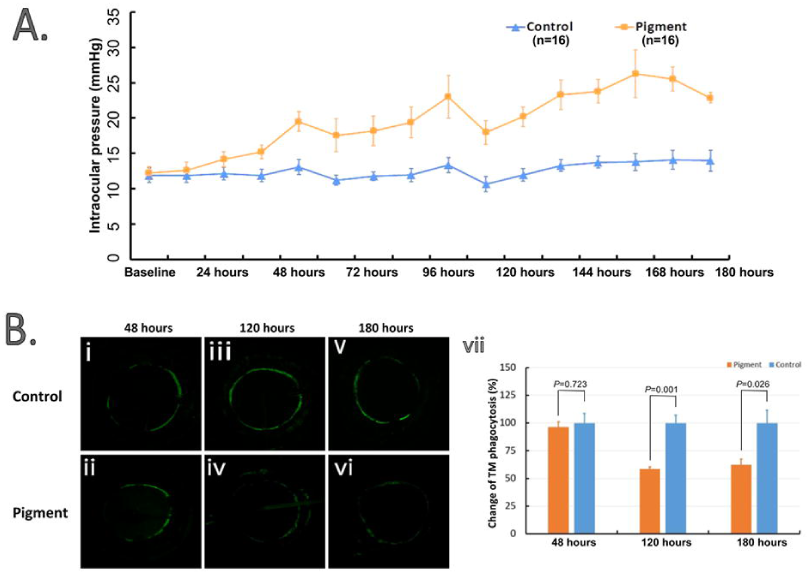
Reduction of intraocular pressure and TM phagocytic activity by pigment dispersion. Baseline IOPs in the pigment group (n=16) and the control (n=16) are comparable (12.2 ± 0.9 mmHg vs. 11.9 ± 0.9 mmHg, *P*=0.82). Pigment caused a significant IOP elevation at 48 hours and onward (all *P*<0.05) while the IOP in the control group showed no significant difference to baseline at any time point (all *P*>0.05) when compared to the baseline (**Fig. 2 A**). TM phagocytosis was visualized in situ. The mean fluorescence intensity in the TM region was quantified by NIH ImageJ. TM phagocytosis in the pigment group was comparable to that of the control at 48 hours (*P*=0.723) (**Fig. 2 B i-ii**) but showed sharp decreases at 120 hours (**Fig. 2 B iii-iv**) and 180 hours (*P*=0.001 and *P*=0.026, respectively) (**Fig. 2 B v-vi**).

## Discussion

Phagocytosis is a defining feature of TM cells^18^ and central to the pathogenesis of several types of secondary glaucoma characterized by particulate matter that include pigment, erythrocytes and ghost cells, inflammatory cells, photoreceptor outer segments, lens or pseudoexfoliation material.^4,19,20^ Although TM phagocytosis can remove particles from the aqueous humor,^21^ the direct and short - term effects of phagocytosis on outflow regulation remain poorly understood.^2^ In this study, we measured IOP and TM phagocytic activity in the presence of pigment granules at different time points and found that IOP was significantly elevated as early as 48 hours after exposure to pigment granules. This was contrasted by a phagocytic activity in P that was not different from C before the decrease at 120 and 180 hours. A worsening decline of TM phagocytosis at 120 and 180 hours did not result in further increase in IOP. This suggests that reduction in phagocytosis is a downstream and secondary effect of actin cytoskeletal reorganization.

Pigment treatment has previously been shown to cause ocular hypertension in part by reorganizing the TM actin cytoskeleton and not by physical obstruction of the outflow tract.^8,22^ We have recently reported that long, thick, and continuous TM actin bundles emerge as early as 24 hours after pigment exposure^4^ and replicate this observation in the present study. Histological characteristics of pigment dispersion in porcine eyes matched those seen in samples from pigmentary glaucoma patients^22–24^ showing that pigment particles were taken up by TM cells.

In summary, the results indicate that the IOP elevation caused by pigment dispersion is not the direct result of a physical obstruction of outflow or a chronically overwhelmed phagocytosis. The reduction in phagocytosis considerably lags the evolving hypertension supporting the notion that these cytoskeletal changes occur early on and are separate from the impact of pigment on canonical phagocytosis pathways.^4^

## Abbreviations

TM: trabecular meshwork
ECM: extracellular matrix
IOP: intraocular pressure
H&E: hematoxylin and eosin

